# Tumor Stroma Content Regulates Penetration and Efficacy of Tumor-targeting Bacteria

**DOI:** 10.1101/2024.03.29.587035

**Authors:** Y. Zhan, B. Burkel, E. J. Leaman, S. M. Ponik, B. Behkam

**Affiliations:** Department of Mechanical Engineering, Virginia Tech, Blacksburg, VA, USA; Department of Cell and Regenerative Biology, University of Wisconsin-Madison, Madison, WI, USA

## Abstract

Bacteria-based cancer therapy (BBCT) strains grow selectively in primary tumors and metastases, colonize solid tumors independent of genetics, and kill cells resistant to standard molecular therapy. Clinical trials of BBCT in solid tumors have not reported any survival advantage yet, partly due to the limited bacterial colonization. Collagen, abundant in primary and metastatic solid tumors, has a well-known role in hindering intratumoral penetration of therapeutics. Nevertheless, the effect of collagen content on the intratumoral penetration and antitumor efficacy of BBCT is rarely unexplored. We hypothesized that the presence of collagen limits the penetration and, thereby, the antitumor effects of tumor-selective *Salmonella*. Typhimurium VNP20009 cheY^+^. We tested our hypothesis in low and high collagen content tumor spheroid models of triple-negative murine breast cancer. We found that high collagen content significantly hinders bacteria transport in tumors, reducing bacteria penetration and distribution by ∼7-fold. The higher penetration of bacteria in low collagen-content tumors led to an overwhelming antitumor effect (∼73% increase in cell death), whereas only a 28% increase in cell death was seen in the high collagen-content tumors. Our mathematical modeling of intratumoral bacterial colonization delineates the role of growth and diffusivity, suggesting an order of magnitude lower diffusivity in the high collagen-content tumors dominates the observed outcomes. Finally, our single-cell resolution analysis reveals a strong spatial correlation between bacterial spatial localization and collagen content, further corroborating that collagen acts as a barrier to bacterial penetration despite *S*. Typhimurium VNP20009 cheY^+^ motility. Understanding the effect of collagen on BBCT performance could lead to engineering more efficacious BBCT strains capable of overcoming this barrier to colonization of primary tumors and metastases.

## Introduction

Bacteria-based cancer therapy (BBCT) has garnered significant interest in recent years (for a comprehensive review, see refs (1, 2)). In preclinical studies, BBCT is shown to selectively grow in primary tumors and metastases (3), colonize solid tumors independent of genetics (4), and kill cells that are resistant to standard molecular therapy (5). However, clinical trials of BBCT have not reported any survival advantage, partly due to the limited bacterial colonization and penetration of cm-sized human solid tumors (2). Self-propulsion and external driving forces, such as magnetic field gradients, have been shown to improve BBCT penetration in solid tumors (6, 7). For instance, we recently demonstrated that the clinically safe (8, 9) and tumor-selective *Salmonella enterica* serovar Typhimurium VNP20009 are effective nano-cargo transporters, enhancing retention and distribution of nanomedicine in low extracellular matrix (ECM) content breast and colon cancer tumors by up to 100-fold (7). However, *S*. Typhimurium VNP20009 poorly penetrated the ECM-rich glioma tumors (7).

Indeed, increased ECM deposition in solid tumors is a prognostic indicator of poor patient outcomes (10–20). Excessive expression of collagen, a primary component of ECM, by cancer-associated fibroblasts (CAFs) promotes higher interstitial pressure and solid stress in tumors, collapses blood vessels, and increases hypoxia, limiting the penetration of anticancer therapeutics (21). The fibrotic ECM is shown to the limited distribution of chemotherapeutics, which can cause clinical resistance (22–25); excludes immune cell infiltration (26); promotes immunosuppression and resistance to immune checkpoint blockade (ICB) therapy (26–28). Nevertheless, the effect of ECM content, particularly collagen, on the penetration and distribution of BBCT and its therapeutic efficacy is rarely explored (29, 30).

Based on our prior findings that *S*. Typhimurium VNP20009 colonizes solid tumors through intercellular (i.e., between cells) translocation (7), we hypothesized that the presence of collagen limits bacterial penetration and thereby antitumor effects of *S*. Typhimurium VNP20009, despite its motility. To test our hypothesis, we chose a murine triple-negative breast cancer model due to the abundance of collagen in its primary tumor and metastases (13) (**Figure 1A-C**) and the well-known role of collagen in hindering intratumoral penetration of therapeutics in breast cancer (23, 31). We developed a low ECM-content homotypic tumor spheroid model comprised of the 4T1 triple-negative murine breast cancer cells and a high ECM-content heterotypic multicellular tumor spheroid model comprised of 4T1 cells and mammary cancer-associated fibroblasts CAFs (mCAFs). We first characterized the collagen content of each tumor model. We then evaluated the intratumoral distribution of bacteria and its antitumor effect in each tumor model as a function of time. We found that high collagen content significantly hinders bacteria penetration in heterotypic tumors. Using a set of quantitative metrics, we show that the penetration depth limit in both tumor types is established within the first few hours and does not improve over time. Thus, the temporal enhancement in the intratumoral distribution of bacteria is almost exclusively due to growth. Our mathematical model of bacteria intratumoral colonization delineates the role of growth and diffusivity, suggesting that the significantly higher diffusivity of homotypic tumors dominates the observed colonization outcomes. We also report that the higher penetration of bacteria in homotypic tumors led to an overwhelming antitumor effect. In contrast, the bacteria-induced antitumor effect in heterotypic tumors was limited to the outer surface due to bacteria being limited to the outer surface. Finally, our single-cell resolution analysis revealed a strong spatial correlation between bacterial distribution and high collagen content, further corroborating that collagen acts as a barrier to bacterial penetration. Understanding how the ECM influences the behavior of therapeutic bacteria within tumors could lead to engineering more efficacious BBCT, capable of overcoming the ECM barrier to colonization of primary tumors and metastases.

**Figure 1.**
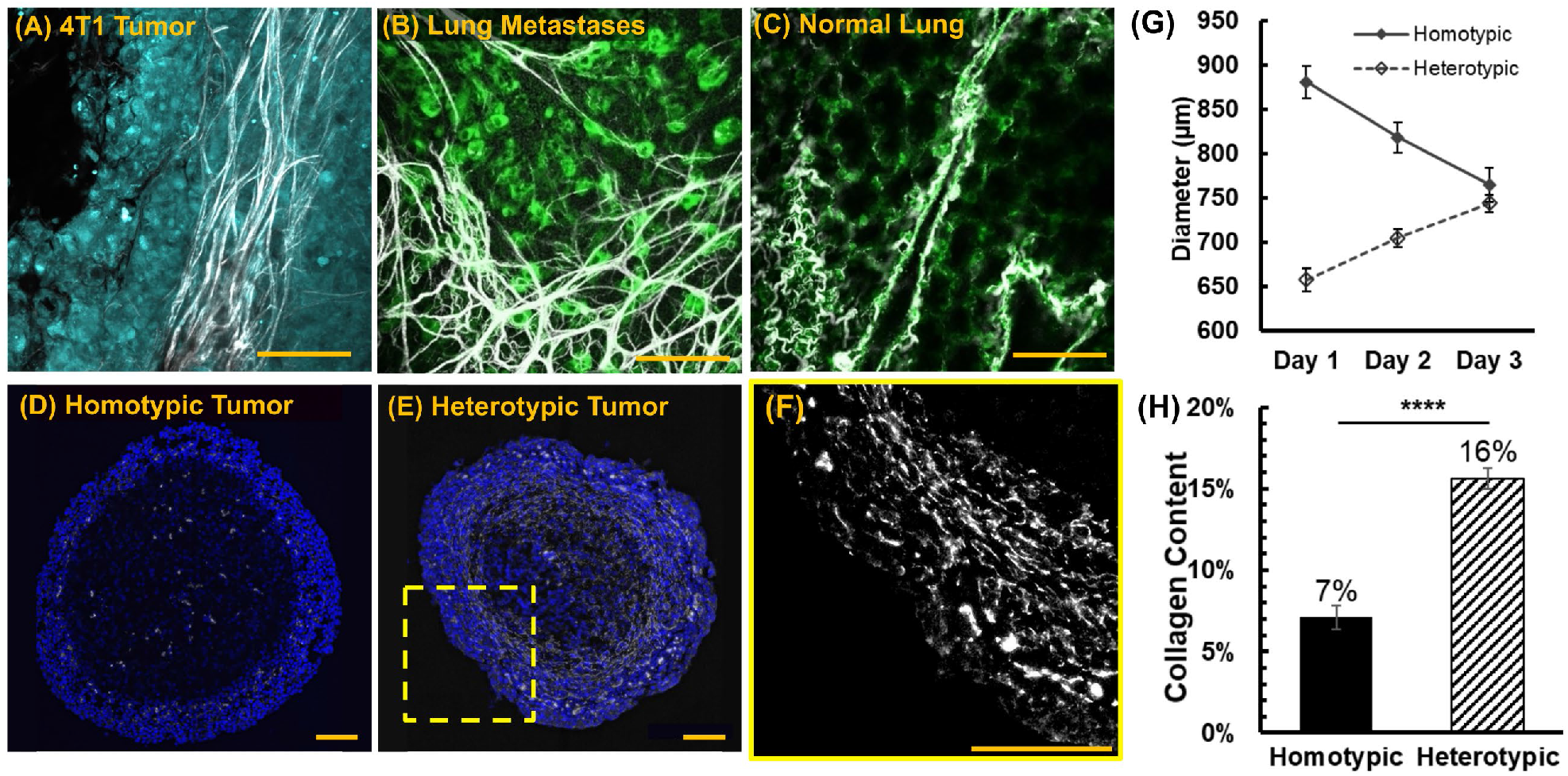
**(A)** Collagen fibers (white) were imaged at the collagen-rich tumor boundary of triple-negative 4T1 tumors *in vivo* and **(B)** in the collagen-rich lung metastasis of tumor-bearing mice. Thick collagen fibers surround metastatic mammary cancer lesions. The collagen content of **(C)** normal murine lung tissue is far less than (B) lung metastatic lesions. Cellular autofluorescence in (B) and (C) is visualized in green. Representative images of collagen type I immunofluorescence staining (white) in homotypic **(D)** and heterotypic **(E)** 4T1 tumor spheroid slices. Cell nuclei in (D)-(E) are stained blue. **(F)** Thick collagen fibers surround the heterotypic tumor, similar to the in vivo primary tumor (A). **(G)** Diameter of tumor spheroid models during the growth phase (prior to inoculation with bacteria, n=9 for each tumor type). **(H)** Collagen content in homotypic and heterotypic tumor slices (n=8 for homotypic tumors and n=10 for heterotypic tumors). In (E) and (G), each data point represents the mean ± standard error. ****p value < 0.0001. All scale bars are 100 μm.

## Materials and Methods

### Mammalian cell culture

Triple-negative murine mammary carcinoma (4T1, ATCC CRL−2539) and murine mammary cancer-associated fibroblasts (mCAFs) (32) were used for the experiments. The 4T1 cells were grown in Roswell Park Memorial Institute (RPMI, Life Technologies, Carlsbad, USA) 1640 medium supplemented with 10% fetal bovine serum (FBS, HyClone). The mCAF cells were grown in modified Dulbecco’s modified Eagle’s medium with high glucose, L-glutamine and sodium pyruvate (DMEM, Corning, Manassas, USA) supplemented with 10% bovine calf serum (BCS, HyClone). Cells were seeded in T-25 flasks for both cell lines with their respective culture medium at 37 °C and 5% CO_2_. For heterotypic tumor experiments, 4T1 cells, cultured in RPMI 1640 with 10% FBS, were adapted to DMEM with 10% BCS for 24 h before making tumor spheroids. All cells were seeded in tissue culture-treated T-25 flasks and incubated in a humidified incubator at 37 °C containing 5% CO_2_.

### Tumor spheroid formation

Once the cell culture in flasks reached a confluency of ∼80% for 4T1 cells or ∼60% for mCAFs, the cells were detached using 1 mL of 0.25% Trypsin-EDTA solution (ATCC, Manassas, USA) followed by 5 min of incubation, and cell densities were estimated using a hemocytometer. To form spheroids of comparable sizes, 60,000 of 4T1 cells (for homotypic tumors) or 34,000 of 4T1 and mCAFs cells at a 1:5 ratio (for heterotypic tumors), respectively, were suspended in 200 μL of DMEM with 10% BCS and seeded in each well of an ultra-low adhesion 96 well plate (Corning Inc., Corning, NY). The well plate was centrifuged at 1000 ×g for 10 min at 22 °C and incubated in a humidified incubator at 37 °C with 5% CO_2_ (33). After two days, the culture medium was replaced with the CO_2_-independent Leibovitz’s L-15 Medium (L-15, Gibco™, New York, USA) supplemented with 10% BCS. The well plate was then incubated in a humidified incubator at 37 °C for one day. The murine 4T1 breast cancer tumor spheroids with and without ECM-depositing mCAFs grew for three days until they reached similar diameters before experiments, as shown in **Figures 1D-G** and **S1**.

### Bacteria culture

The tumor-targeting chemotactic *Salmonella* Typhimurium VNP20009 cheY^*+*^(34) was transformed with a plasmid encoding constitutive expression of a red fluorescent protein (RFP). The plasmid was constructed by inserting the BioBrick part (iGEM Foundation, Cambridge, MA), BBa_J04450, encoding mRFP1 expression in the high copy number pSB1C3 vector with resistance to chloramphenicol. For each experiment, 10 mL MSB medium (34) (1% tryptone, 0.5% yeast extract, 2 mM MgSO_4_, 2 mM CaCl_2_, pH 7.0) in a 125 mL smooth-bottom flask, supplemented with 35 μg/mL of chloramphenicol, was inoculated with a single colony of *Salmonella* Typhimurium VNP20009 cheY^+^, selected from a 1.5% MSB agar plate culture. After overnight incubation at 37°C and 100 rpm, the bacterial culture was diluted 100× in MSB supplemented with chloramphenicol. This diluted culture was grown at 37°C and 100 rpm to establish a new culture in the exponential growth phase. After reaching an optical density at 600 nm (OD_600_) of 1.0, bacterial culture was washed twice at 1700 ×g for 5 min with L-15 medium supplemented with 10% BCS and concentrated to a final OD_600_ of 1.0 in the same medium prior to use in the experiment.

### Bacteria intratumoral penetration assay

Six tumor spheroids of the same type (homotypic or heterotypic) were transferred into a 1.5 mL microcentrifuge tube and incubated with bacteria at OD_600_ = 1.0 (approximately 4.5×10^8^ colony forming units (CFU)) in 1 mL of L-15 medium with 10% BCS. The tubes were rotated on an end-over-end shaker at 15 rpm at 37 ºC to allow for thorough mixing and uniform conditions. After 1 h, tumor spheroids were washed with fresh culture media, incubated within cell culture medium supplemented with 2.5 μg/mL gentamicin sulfate, and returned to the shaker to kill any residual bacteria in the suspension. After 30 min of antibiotic treatment, tumor spheroids were washed with fresh culture media, suspended in 1 mL of L-15 medium with 10% BCS and put back on the shaker at 15 rpm at 37 ºC. Tumor spheroids were harvested after 12, 18, 24, 36, and 48 h total incubation period and gently rinsed with Dulbecco’s phosphate-buffered saline (DPBS, Gibco) to wash away the bacteria loosely associated on the surface of the tumor spheroids.

### Immunohistochemistry

The tumor spheroids were transferred into cryomolds coated with a thin frozen layer of 1:1 (v/v) mixture of optimal cutting temperature (OCT) compounds and 60% w/v sucrose in deionized water and then filled with the same mixture. The tumor spheroids were frozen at −20 °C overnight before sectioning. A cryotome was used to section the tumor spheroids into 10 μm thick slices, which were transferred onto a SuperFrost Plus glass slide (Fisherbrand). For evaluation of intratumoral penetration of bacteria, each tumor slice was stained with a droplet of nucleus stain NucBlue (Thermo Fisher Scientific, Waltham, MA) with the concentration of 2 drops/ml in DPBS for 15 min at room temperature and covered with a coverslip. For evaluation of the bacteria antitumor effect, each tumor spheroid slice was thawed and fixed with 4% paraformaldehyde (PFA) for 20 minutes at room temperature. The slides were then rinsed with PBS (3×) and then stained with an *in situ cell death detection* (FITC-TUNEL) staining kit (Millipore), an AlexaFluor-647 anti-collagen antibody (NovusBio, 2749AF647), and DAPI.

### Image acquisition and processing

For bacterial intratumoral penetration evaluation, fluorescence microscopy images of DAPI-stained tumor slices were collected using a Zeiss AxioObserver Z1 inverted microscope equipped with an AxioCam mRM camera and a 40× objective. Tumor cells were visualized using 365 nm excitation wavelength with a 445/50 band-pass DAPI (49) emission filter. Bacteria were visualized using 550/25 excitation wavelength with a 605/70 band-pass DsRed (43 HE) emission filter. Bacteria intratumoral distribution was quantified using our previously developed imaging processing algorithm in MATLAB (7). Imaged slices were binned into rings of uniform 10 μm width, and bacteria numbers were determined from the fluorescence area and the average size of a single bacteria. To evaluate the spatial distribution of bacteria, a minimum of 3 independent experiments and 3 tumors for each experiment category were used. Up to 3 slices from each tumor were taken near the spheroid’s great circle. For evaluation of collagen content and bacteria antitumor effect, the multiplexed immunohistochemistry (IHC) slides were imaged using a Leica Thunder imaging system with a DMi8 microscope base, an 8-line LED, and a DFC9000 GT scMOS camera. The following excitation and emission filter combinations were: DAPI was visualized with a 391/32 excitation filter and a 435/30 emission filter, the FITC conjugated TUNEL stain was visualized with a 479/33 excitation filter and 519/25 emission filter, the bacteria were visualized using 554/24 excitation filter and a 594/32 emission filter, and collagen was visualized using a 638/31 excitation filter and a 695/58 emission filter. Tiled z-stacks of the entire spheroid slices were collected with an HC PLAN APO 40×/1.10 WATER objective with the z-stacks collected at Nyquist in each condition. The illumination levels were set to maximize each channel’s dynamic range and held constant across conditions. The tiled stacks were then merged and deconvolved with the Large Volume Computational Clearing module within the LAS X software and exported as TIFFs for additional analysis within ImageJ.

### Metrics for quantitation of bacteria intratumoral transport performance

Our previously developed penetration index (PI), colonization index (CI), and distribution index (DI) (7) were used to quantitate bacteria intratumoral distribution and delineate the role of bacterial growth from penetration in the observed colonization trends. PI is defined as 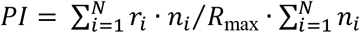, where *r*_*i*_ is the radial penetration distance of the *i*^th^ bin from the spheroid surface, *n*_*i*_ is the number of the bacteria within the *i*^th^ bin, and *R*_max_ is the maximum penetration distance to the center of the spheroid. The PI value ranges between 0 (for bacteria on the surface of the spheroid) and 1 (for bacteria at the center of the spheroid). CI is defined as 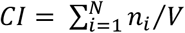, where *V* is the volume of the tumor slice. It measures the volumetric density of bacteria in units of CFU/μm^3^. Finally, DI is a combinatorial metric *DI* = *PI × CI*, representing the extent of bacteria distribution in the tumor.

### Cell viability assay

To examine the bacteria effect on tumor cell viability, live/dead assays at different timepoints (12, 18, 24, 36, and 48 h) in the presence and absence of bacteria (control) were performed. Tumor spheroids were digested into single cell suspensions using 0.028 Wünsch units/ml of Liberase TM Research Grade (Roche, Mannheim, Germany) and 20 μg/ml of DNase I in L-15 with 10 % CS for 1 hour at 37 °C and 250 RPM on an orbital shaker. Agitation was facilitated by pipetting up and down every 10 minutes. The cells were suspended in L-15 with 10 % CS, NucBlue® Live reagent (2 drops/ml) and NucGreen® Dead reagent (2 drops/mL) in a well plate. The fluorescent microscopy images of the tumor slice were taken using a Zeiss AxioObserver Z1 inverted microscope equipped with an AxioCam mRM camera and a 10× objective. Cells were counted manually for each image and averaged per tumor spheroid.

### Statistical Analysis

All reported results are based on a minimum of three independent experiments, and results were averaged across 2-3 sections from each tumor, when applicable. All data are presented as means ± standard error. All statistical analyses were performed in OriginPro (OriginLab, Northampton, MA). Pairwise comparisons were conducted using a *t*-test. Data with more than two categories were analyzed using the one-way analysis of variance method followed by Fisher’s least significant difference test. Statistical significance was defined as p-values less than 0.05.

### Mathematical modeling of the bacterial tumor spheroid colonization

Bacterial intratumoral transport was modeled as a diffusive process that would capture intratumoral penetration due to growth and motility. The transport, colonization, and nutrient availability were described using diffusion-reaction equations of the form

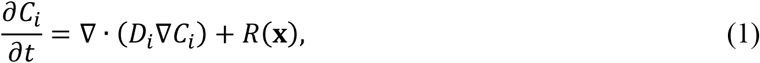

where *C*_*i*_ is the local concentration of either bacteria or nutrients, *D*_*i*_ is the diffusion coefficient of *C*_*i*_, and *R*(**x**) is the first-order reaction terms (**x** = *x, y, z*) that represent growth and media consumption. Assuming symmetry in the angular dimensions, a set of coupled partial differential equations describe normalized intratumoral penetration and colonization of bacteria, which in spherical coordinates are:

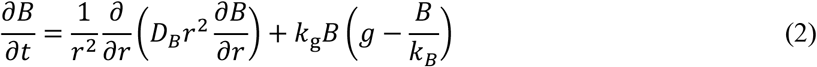

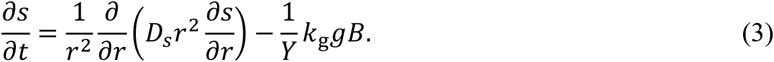

Equations 2 and 3, respectively, govern the transport and growth of bacteria, *B*, and the transport and utilization of substrate (cell nutrient medium) *s*, with respect to radial position *r*. The diffusion coefficients are given by *D*_*i*_, *k*_*g*_ is the maximum growth rate of *B* (*k*_*g*_ = ln(2) /*τ*_min_, where *τ*_min_ represents the doubling time in a nutrient-rich medium), *k*_*B*_ = *k*_*B*_ (*r*) is the local carrying capacity of the tumor (*i*.*e*., the maximum possible concentration of *B*) (35), and *Y* is the bacterial yield (*i*.*e*. the ratio of bacteria produced to amount substrate consumed). The local growth rate is governed by the substrate availability according to Monod growth kinetics,

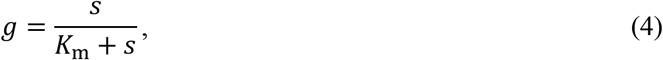

where *s* = *s*(*r*) is the local substrate concentration with respect to radial position *r*, and *K*_m_ is the Monod constant (*i*.*e*., the concentration of *s* when the growth rate of *B* is one-half the maximum rate).

The boundary conditions for all diffusing species at the center of the tumors were taken as

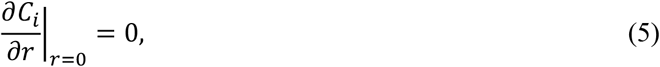

and the boundary conditions at the tumor peripheries were

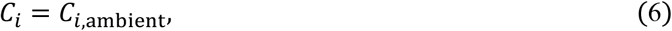

where *C*_*i*,ambient_ is the ambient concentration of *B* or *s*.

### Model Fitting

Each simulation was executed to capture the experimental procedure. The modeled bacterial and substrate concentrations were normalized to their ambient concentrations at the start of experiments. Prior to *t* = 0, tumors were simulated to be in the steady state, saturated with growth medium. At *t* = 0, the bacterial concentration in the surrounding medium (ambient) was set to its normalized value of unity. At *t* = 1 h, we simulated the introduction and effects of the antibiotic supplement by assuming that only 0.05% of bacteria in the ambient medium survived the 30-minute incubation period, while 10% of those residing intratumorally survived. The normalized ambient concentration of cell growth medium was set to 1 following the treatment (at *t* = 1.5 h), simulating replenishment of fresh medium.

The model was fitted to the measured bacterial concentration curves by optimizing the values of *D*_*B*_ and *τ*_min_ using a differential evolution algorithm. Growth parameters *K*_m_, and *Y* were assumed to fall between 0.4 and 0.6 and 0.6 and 0.8, respectively, and were each allowed to vary within these ranges for fitting the data. The constant model parameters *Y* and *D*_*s*_ were estimated at 10 (normalized bacteria/normalized substrate) and 100 μm^2^/s (glucose diffusivity in tumor spheroids (36)), respectively.

## Results and Discussion

### Effect of ECM on bacteria intratumoral penetration and colonization

We used homotypic 4T1 tumor spheroids (**Figure 1D**) and heterotypic tumor spheroids (**Figure 1E-F**), comprising 4T1 cancer cells and mCAFs, to assess how changes in the tumor ECM content affects intratumoral transport and colonization of bacteria. As shown in Figures **1A** and **1F**, our heterotypic 3D model recapitulates matrix content and organization of the *in vivo* tumors. Using immunofluorescence staining of collagen Type 1, we show that the collagen content of heterotypic 4T1/mCAF spheroids is 2.3× higher (*p <* 0.0001) than the 4T1 homotypic tumors (**Figure 1H**).

The homotypic and heterotypic tumor spheroids were incubated with RFP-expressing *S*. Typhimurium VNP20009 cheY^+^ (hereafter referred to as bacteria) for 12 h, 18 h, 24 h, 36 h, and 48 h. To image and analyze bacteria penetration and colonization, tumor spheroids were sliced into 10 μm-thick sections and stained with DAPI (see Methods). Three to five sections with the largest cross-sectional area were imaged using fluorescence microscopy. In homotypic tumors, bacteria were observed to have penetrated to the tumor core within the first 12 h (**Figures 2A-i** and **2C**). Bacteria-colonized regions grew larger over time and began to affect the integrity of the tumors. At 36 h of co-incubation with bacteria, some of the homotypic tumors disintegrated (Figures 2A-iv), while at 48 h, all homotypic tumors completely disintegrated into fragments, as shown in Figure 2A-v (as a result, bacterial distribution at 48 h are not included in Figure 2C). In contrast, bacterial penetration in the heterotypic tumors was quite limited within the first 12 h (**Figure 2B-i**) and did not exceed 25% of the radial distance from the tumor’s surface (**Figure 2D**). Over the 48 h experiment period, the size of the bacteria colonized regions grew, but bacteria remained primarily limited to <50% of the radial distance and did not reach the tumor core (**Figure 2B** and **2D**). Heterotypic spheroids maintained their integrity throughout the 48 h experiment period, presumably due to the higher ECM content and the severely limited bacteria penetration.

**Figure 2:**
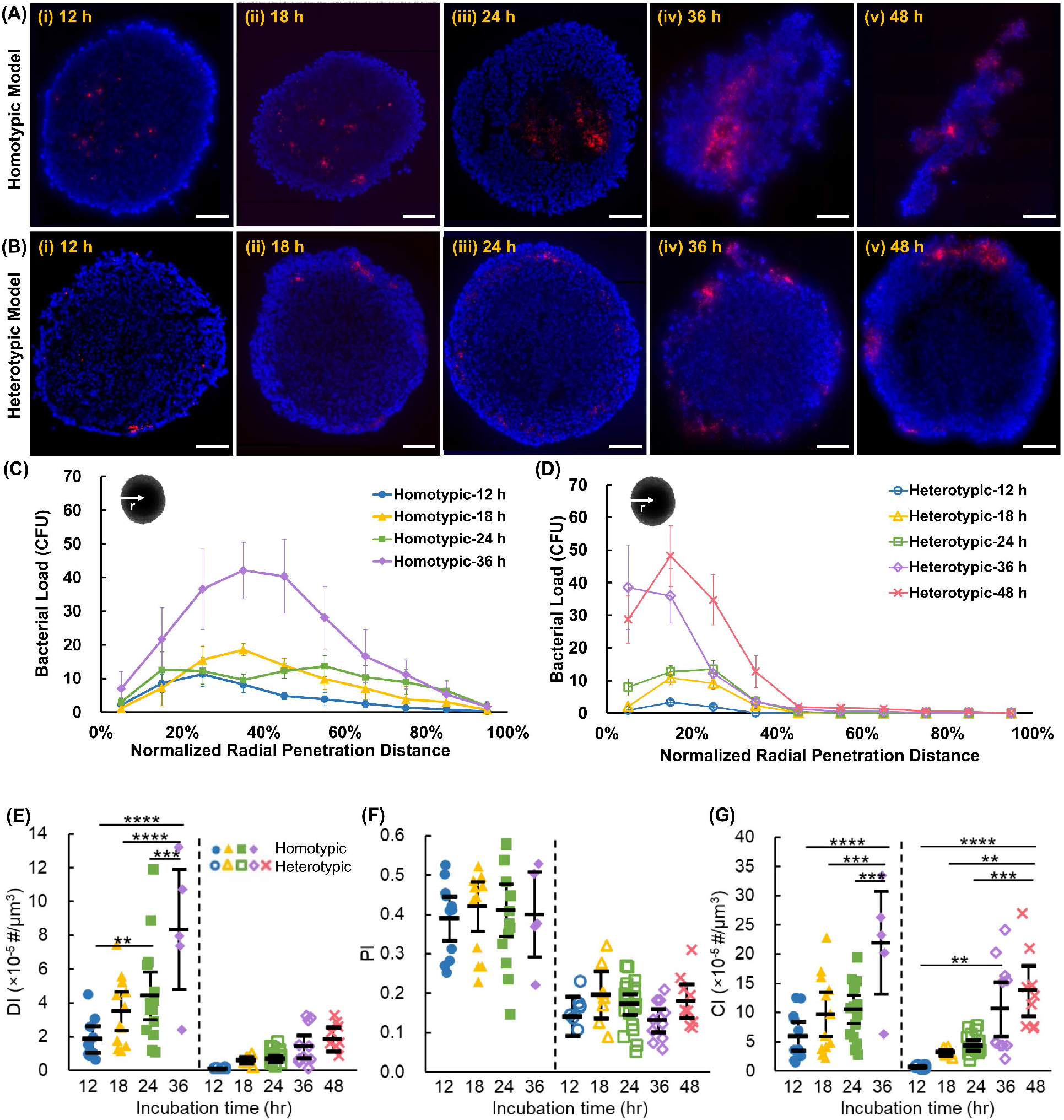
Representative composite fluorescence microscopy images of **(A)** homotypic and **(B)** heterotypic tumor sections at 12, 18, 24, 36, and 48 h after incubation with bacteria. Tumor cell nuclei are stained in blue and bacteria express RFP (red). Bacterial concentration versus normalized radial penetration distance from the tumor surface in **(C)** homotypic and **(D)** heterotypic tumor models. Each data point represents the mean ± standard error. Quantitative indices of **(E)** distribution index (DI), **(F)** penetration index (PI), and **(G)** colonization index (CI) indicate bacteria intratumoral transport performance in homotypic and heterotypic tumors. The black bars indicate means and 95% confidence intervals. \*\**p*-value<0.01, *** *p*-value<0.001, \*\*\*\**p*-value<0.0001. In (C)-(G), n = n = 11, 13, 16, and 5 independent tumors at 12, 18, 24, and 36 h for homotypic tumors and n = 8, 7, 20, 11, and 9 independent tumors at 12, 18, 24, 36, and 48 h for heterotypic tumor spheroids, respectively. Experimental results were averaged across 3-5 sections from each tumor. All scale bars are 100 μm.

We previously developed three metrics, penetration index (PI), colonization index (CI), and distribution index (DI) (see Methods), to delineate the role of penetration and colonization (growth) in the intratumoral distribution of therapeutic agents (7). Overall, the DI of homotypic tumors is significantly higher than that of the heterotypic tumors at each incubation time (**Figure 2E**). Consistent with the data shown in Figures 2C-D, the DI values increased in both tumor types over time; however, the rate of increase in homotypic tumors was significantly higher, culminating in a 5.9-fold difference in the DI values of the two tumor types at the 36 h timepoint. Another striking observation was that after 48 h, the heterotypic tumors reached DI values similar to those of homotypic tumors after only 12 h. To delineate the effect of growth from penetration, we investigated the contribution of PI and CI to the observed DI trends. Notably, the PI value for homotypic tumors (0.41±0.01) was 2.4-fold higher than the PI value for heterotypic tumors (0.17±0.02), suggesting that the significantly higher collagen content in heterotypic tumors (Figure 1H) inhibits the intratumoral transportation of the bacteria. Interestingly, there was no statistically significant change in the PI values over time independent of the tumor type (**Figure 2F**). Although the role of collagen content increase in intratumoral penetration of self-propelling bacteria was not studied prior to this work, our findings are consistent with prior works reporting reduced penetration of other therapeutics in high collagen content tumors(22–27, 31, 37, 38). Our results show that *Salmonella* Typhimurium VNP20009 cheY^*+*^ motility is insufficient to overcome this biological barrier in 4T1 murine breast cancer tumors. Unlike the PI trends, the CI in both homotypic and heterotypic tumors increased significantly, with increased incubation time (**Figure 2G**). An increase in CI indicates increased colonization, which is expected due to bacterial growth in the nutrient-rich environment of the tumors. Nonetheless, it should be noted that the homotypic tumors had significantly higher CI values than heterotypic tumors at all timepoints. For instance, the CI of the homotypic tumors (21.9±10.0) was 2.1× higher than that of heterotypic tumors (10.6±7.8) at the 36 h timepoint. We attribute this difference to the higher initial entrapment of bacteria in the homotypic tumors, which leads to more significant differences over time due to the exponential growth combined with the denser microenvironment in the heterotypic tumor, which may present resistance to bacterial fission and leads to reduced growth. Altogether, our quantitative analyses demonstrate that the high ECM content of the physiologically relevant tumor models limits bacterial penetration and growth despite the self-propulsion capability of tumor-targeting *S*. Typhimurium.

### Mathematical model of bacterial colonization of tumors

We developed a mathematical model to delineate the effect of increased collagen content on bacteria growth and intratumoral transport. The primary goal of this mathematical model was to produce quantitative transport (diffusion coefficient or diffusivity) and growth (doubling time) metrics for comparing bacteria performance in the two tumor models. We assumed bacterial transport within the tumors could be approximated as a Fickian diffusion process and captured growth both inside and outside (in the surrounding suspension medium) the tumors using Monod kinetics, with the cell culture medium as the substrate source (see Methods).

As shown in **Figure 3A**, our mathematical model captured the bacteria distribution in homotypic and heterotypic tumors across all timepoints. The fitted diffusivity and doubling time (*i*.*e*., the minimum doubling time for an ideal substrate) for representative tumor slices are depicted on the corresponding graph. Increased spatial heterogeneity in tumor colonization at later timepoints led to slightly increased differences between the experiment and the fitted in homotypic tumors since the model assumes uniformity in the angular dimension. Aggregated model fitting data showed that the average diffusivity of bacteria in homotypic tumors was about an order of magnitude higher at each incubation timepoint than that of the bacteria in heterotypic tumors (**Figure 3B**). Expectedly, there was no significant difference in effective bacteria diffusivity as a function of time in either of the tumors. The high standard error in homotypic tumor diffusivity at 36 h is due to extensive colonization of bacteria and significant tumor deformation, which leads to considerable differences between replicates (**Figure S2**). Nonetheless, the fitted diffusivity values are in agreement with our experimental observations. The heterotypic tumors maintained good integrity even after long incubation periods with bacteria and contained abundant collagen type I, which is suggested to limit bacterial penetration in tumors (7). In contrast, homotypic tumors often contained voids in tumor sections; some disintegrated within 24 h, and they contained little collagen. In agreement, bacteria penetrated and colonized homotypic tumors to a greater extent, reflected in the significantly larger fitted diffusivities.

**Figure 3:**
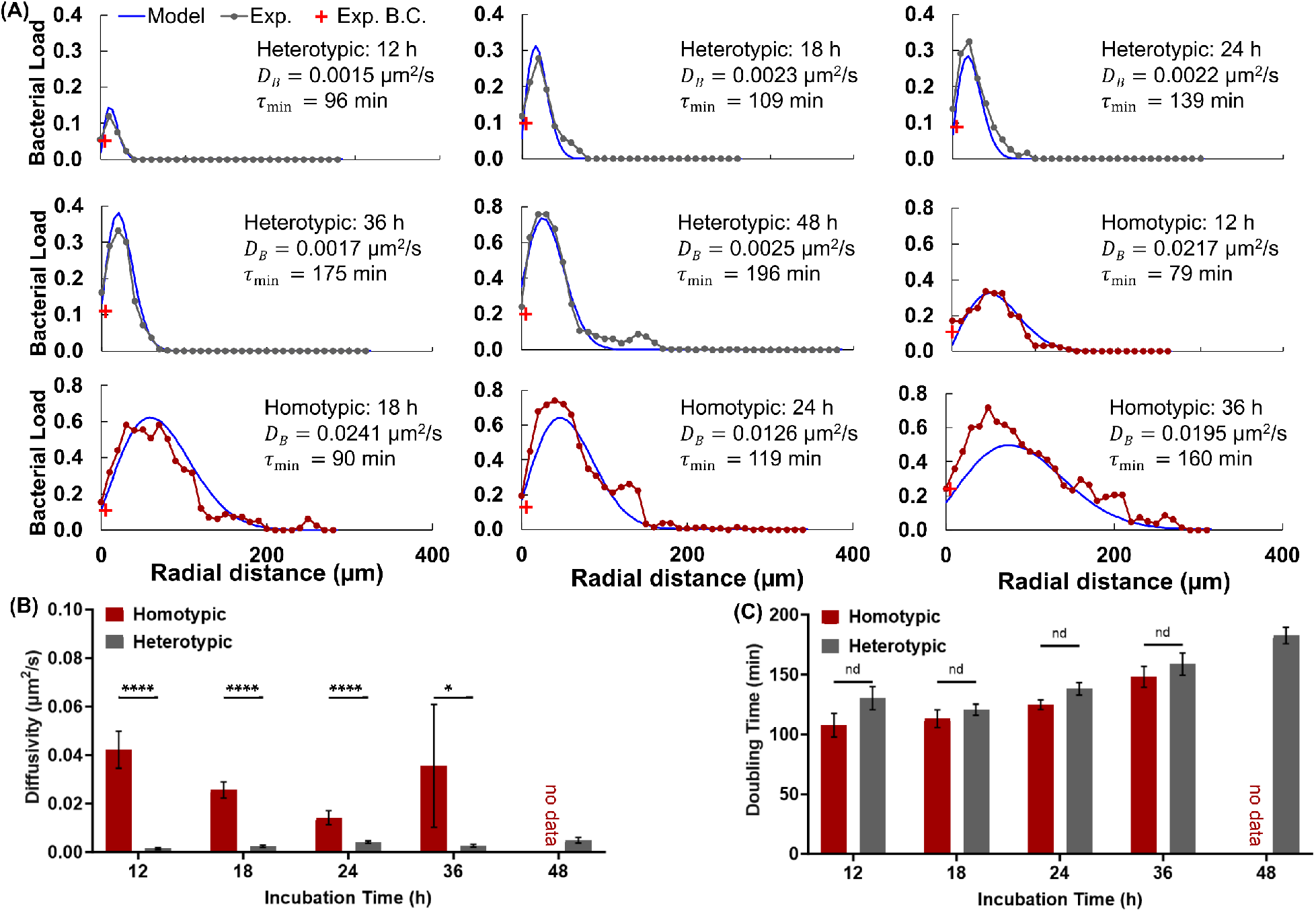
Mathematical Model Fitting Results to Experimental Intratumoral Bacteria Concentration Curves. **(A)** Representative fitting results (blue) and the corresponding experimental data (black), and bacteria concentration in the ambient media (red) for homotypic and heterotypic tumors at each point in time. The incubation time, fitted diffusivity and doubling time values are shown on each plot. **(B)** Average best-fit diffusivity and **(C)** average best-fit doubling time values at each incubation time for homotypic and heterotypic tumors. Error bars in (B) and (C) represent ±S.E.M. For homotypic tumors, *n* = 10, 13, 14, from 4 independent tumors, and for heterotypic tumors, *n* = 8, 7, 22, 11, from 6 independent tumors at 12, 18, 24, 36, and 48 h, respectively. \*\*\*\**p*<0.0001 and \**p*<0.05.

In contrast to the diffusivity trends, the model fitting showed only a slight increase in doubling time in heterotypic tumors, and the differences between the two tumor types were not statistically significant (**Figure 3C**). The shorter average doubling times for the homotypic tumors may be attributed to the lower physical constraints (*i*.*e*., collagen) and stiffness in this tumor type.

### Effect of bacterial intratumoral distribution on tumor cell death

We next inquired how the differences in bacteria penetration in homotypic and heterotypic tumors affect tumor cell death. We digested the control and bacteria-treated tumors at 12, 18, 24, 36, and 48 h timepoints to prepare tumor cell suspensions and assessed cell viability using a cell viability imaging kit (see Methods). We measured >80% tumor cell viability in both homotypic and heterotypic control tumors at all timepoints (**Figure S3A**). The dead cell fraction was measured at 4% at 12 h, increased to 14-17% at the next measurement point of 18 h, and remained unchanged for the remainder of the 48 h experiment period. The presence of dead cells in the control tumors is attributed to the formation of a necrotic core due to the size of tumor spheroid models. There was no statistically significant difference in the fraction of dead cells between the homotypic and heterotypic control tumors at each timepoint (Figure S3A).

Comparatively, bacteria-treated tumors had a higher fraction of dead cells at all timepoints. Moreover, there was a steady increase in the number of dead cells with time (**Figure S3B**). Pairwise comparisons of homotypic and heterotypic tumor data revealed statistically significant differences at most timepoints. Strikingly, the homotypic tumors had only 10% viable cells (90% dead cells) after 48 h, whereas 58% of the cells were viable in heterotypic tumors. To distinguish the effect of bacteria on tumor cell death from the effect of hypoxia, we examined the differential cell death percentage between the bacteria-treated and control tumors (**Figure 4A**). The antitumor effect of tumor-selective *S*. Typhimurium VNP 20009 in homotypic tumors progressively increases in the homotypic tumors, leading to near-complete tumor disintegration and a 73% increase in tumor cell death, compared to the control homotypic tumors. In contrast, bacteria only had a modest antitumor effect on heterotypic tumors, causing a maximum of 29% increase in tumor cell death at 48 h, compared to the control heterotypic tumors.

**Figure 4:**
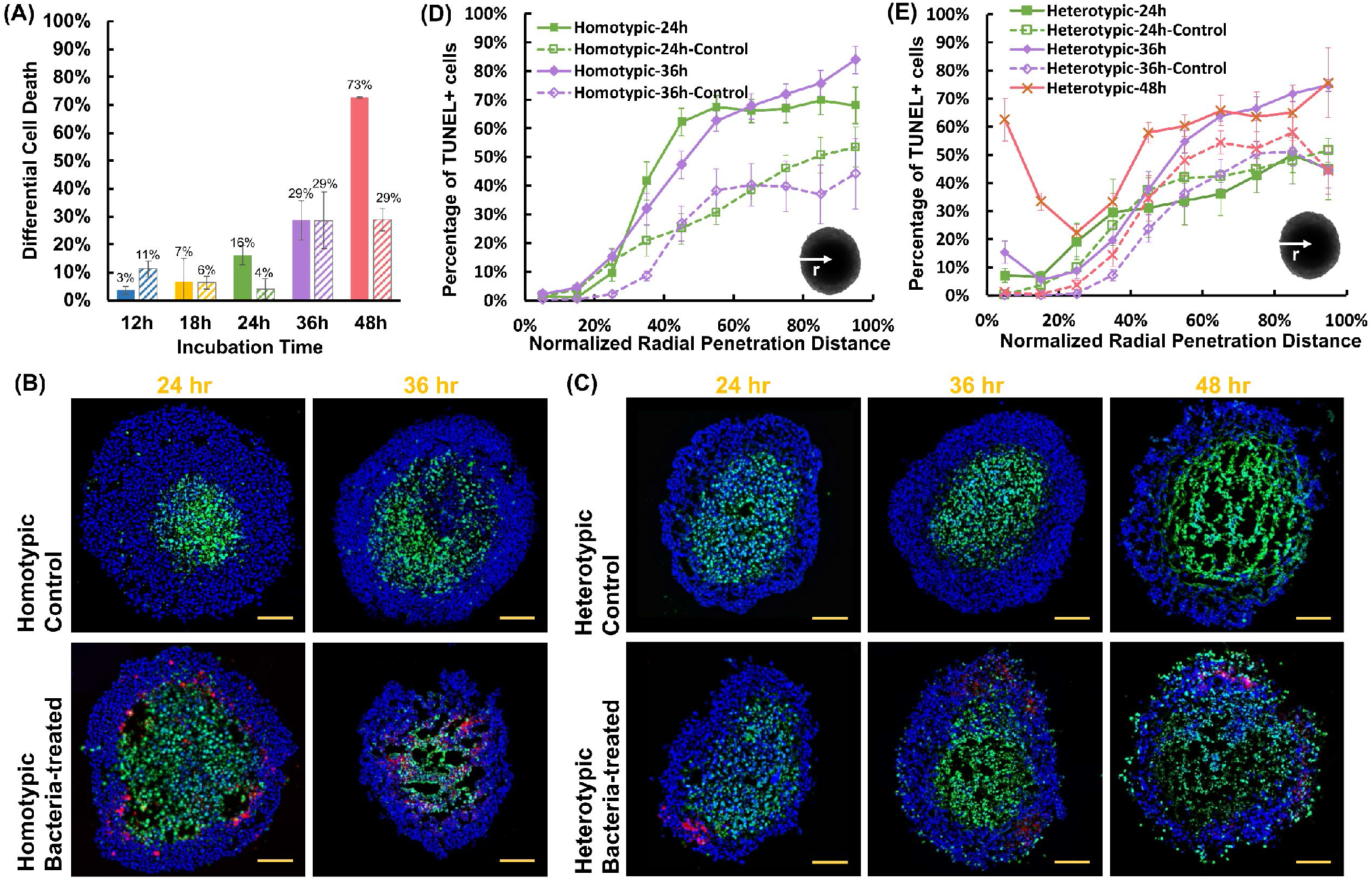
**(A)** The difference in the percentage of TUNEL-positive cells between control and bacteria-treated homotypic and heterotypic tumor spheroids. For control homotypic tumor spheroids, n = 5, 3, 3, 3 from 3 independent tumors, and for heterotypic tumor spheroids, n = 4, 3, 3, 3 from 3 independent tumors at 12, 18, 24, 36, and 48 h, respectively. For bacteria-treated homotypic tumor spheroids, n = 10, 3, 8, 5 from 3 independent tumors, and for bacteria-treated heterotypic tumor spheroids, n = 7, 6, 4, 3 from 6 independent tumors at 12, 18, 24, 36, and 48 h, respectively. All results are from the collection of at least 100 total cells from each independent tumor spheroid. Representative confocal microscopy images of control and bacteria-treated homotypic **(B)** and heterotypic **(C)** tumor sections for TUNEL assay. In all images, tumor cells are stained in blue (DAPI), TUNEL-positive cells in green and bacteria in red. All scale bars are 100 μm. Distribution of TUNEL-positive cells from the surface to the center of **(D)** homotypic and **(E)** heterotypic tumors at each timepoint. Colored curves for each case show the averages at each timepoint. For homotypic tumors, n = 3, 3 (control) and n = 11, 8 (bacteria-treated), at 24, and 36 h from 3 independent experiments, and for heterotypic tumors n = 5, 5, 6 (control) and n = 3, 5, 6 (with bacteria), at 24, 36, and 48 h from 3 independent experiments, respectively. In all plots, each data point represents the mean ± standard error. Statistical analyses indicate pairwise comparison between homotypic and heterotypic tumors at each timepoint.

We next inquired about the differences in the spatial distribution of apoptotic cells in homotypic and heterotypic tumors and any potential correlation with bacterial colonization sites. We used terminal deoxynucleotidyl transferase-mediated dUTP nick end labeling (TUNEL) assay to detect DNA fragmentation caused by the activation of the apoptotic signaling cascades in control and bacteria-treated tumor slices using confocal microscopy (**Figure 4B-C**). Guided by the viability assay results, we limited our spatial analysis to 24, 36, and 48 h timepoints. Analysis of homotypic tumors at 48 h timepoint was not possible due to the disintegration of the tumors and near-complete tumor cell death at this timepoint (Figure S3B). Both types of control tumors displayed comparable trends in the percentage of TUNEL-positive cells during the entire incubation period (**Figure S4A**). The portion of TUNEL-positive cells peaked at ∼50-55% at >70% radial distance from the tumor surface and remained largely unchanged towards the center. This observation suggests that the increased collagen content does not significantly alter the diffusion of oxygen and nutrients into the heterotypic tumor models.

Comparatively, bacteria treatment of homotypic tumors led to an up to 37% rise in the number of TUNEL-positive cells at the 24 h incubation period, compared with the control (**Figure 4D**). The maximum rise occurred in the intermediate radius range (40-65%). At 36 h, a similar increase of up to 40% in the percentage of TUNEL-positive cells was observed. However, the increase spanned the larger radius range of 35%-100%. It should be noted that the profound antitumor effect of the bacteria on homotypic tumors caused the development of “hollow” regions in the tumor core (Figure 4B), precluding the inclusion of dead cells that were lost during the sample preparation process in the analysis. In the heterotypic model, bacteria treatment did not cause a significant change (<10%) in the TUNEL-positive cell percentage at 24 h (**Figure 4E**). At 36 h, a modest increase of up to 21% at the core was measured. Interestingly, TUNEL-positive cells began to emerge in the peripheral region of the heterotypic tumors at 36 h and increased with time (**Figure 4C** and **4E**). By 48 h, over 60% of the cells in the peripheral region of the heterotypic tumors were TUNEL-positive. This higher quantity of TUNEL-positive cells appears consistent with the higher bacterial density in the peripheral regions (viable rim) of the heterotypic tumors, wherein the bacterial distribution is confined to the outermost 30% of the radius (Figure 2A). These results suggest that the superficial tumor colonization by bacteria contributes to the increased occurrence of apoptotic cells in the viable rim; however, due to the overall lower colonization of heterotypic tumors (Figure 2D), the bacteria antitumor effect is smaller (Figure 4A).

Altogether, our results show that in the easy-to-penetrate homotypic tumor model (Figure 2A, C), bacteria effectively penetrate the tumor and induce an increase in the number of TUNEL-positive cells (Figure 4D). In contrast, in the high collagen content hard-to-penetrate heterotypic tumors, bacterial colonization is limited to the peripheral region of the tumors (Figure 2B, D), leading to the presence of TUNEL-positive cells in these regions, with no significant change in the cell viability in the core (Figure 4E).

### Spatial distribution of TUNEL-positive cells, bacteria, and collagen in the heterotypic tumors

Given the spatial patterns for bacteria and TUNEL-positive cell distribution in heterotypic tumors (Figures 2D and 4E), we inquired if the regions of high collagen density in these tumors present a barrier to bacterial penetration and lead to peripheral accumulation of bacteria in the viable rim. We then asked if such localized colonization could induce regional tumor cell death. To answer these questions, we carried out high-resolution confocal microscopy of heterotypic tumor slices (**Figure 5A-B**). We then defined regions of interest (boundaries of representative ROIs are shown in orange, **Figures 5A.i** and **S5**), which encompassed high collagen density zones (bright white spots) and studied the co-localization of bacteria and collagen and its relation to the regional distribution of TUNEL-positive cells. We observed that the locations of maximal bacterial accumulation coincide with the peak collagen content (**Figures 5C** and **S5**). Examination at higher spatial resolution (**Figure 5A.ii**) shows an even stronger correlation between the bacteria location and collagen distribution (**Figure 5D**). Our findings suggest that collagen indeed presents a barrier to bacterial penetration. Since *S*. Typhimurium VNP 20009 does not invade 4T1 cells (7), bacteria penetration is impeded in regions of high collagen density. Bacteria growth in these dense regions locally increases bacteria number over time (Figure 2B) but they cannot overcome the collagen barrier to penetrate further.

**Figure 5:**
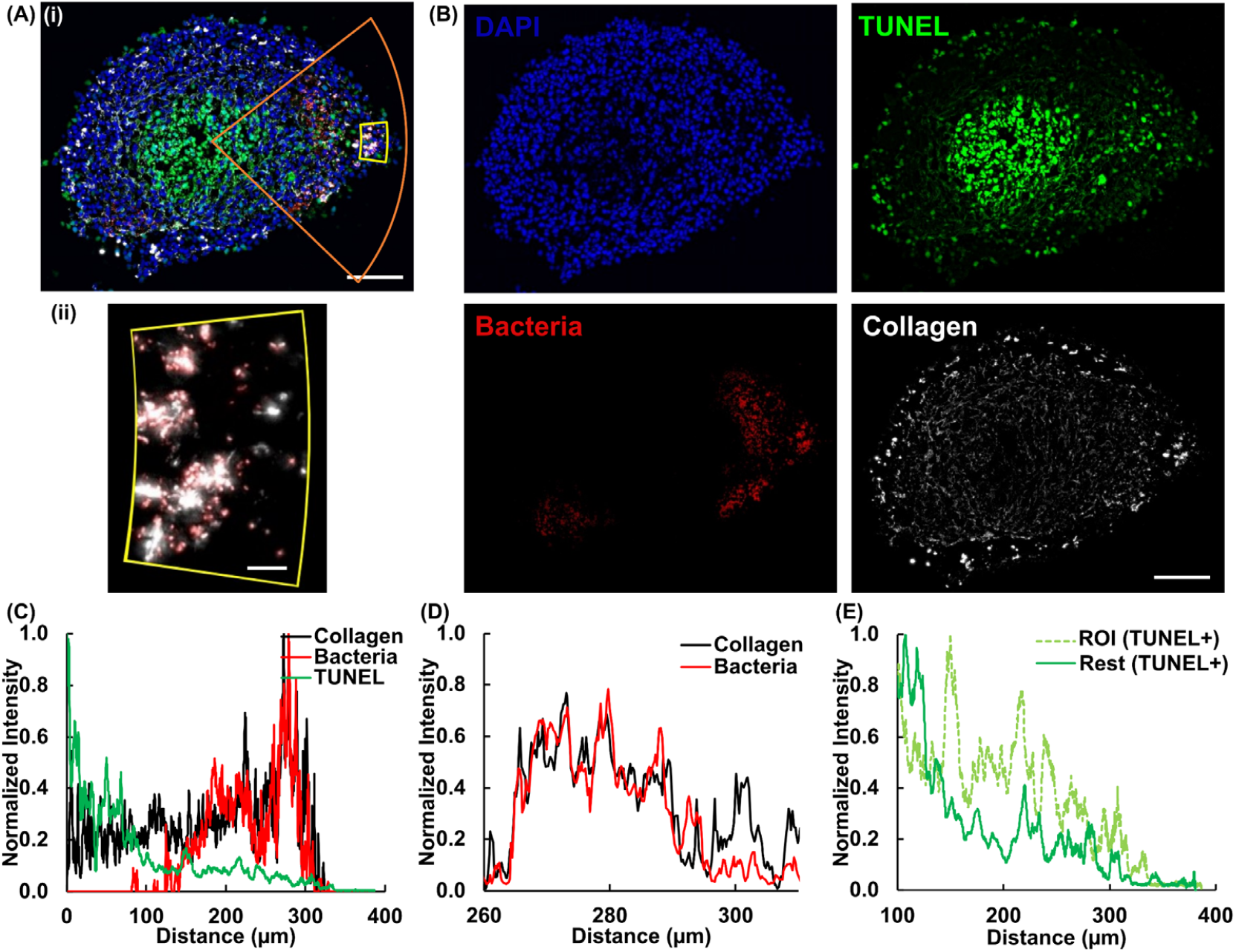
**(A)** Composite fluorescence microscopy image of heterotypic tumor slice 36 h after colonization (i) and composite fluorescence microscopy image of a zoomed-in region (yellow box in 4A) showing collagen (white) and bacteria (red). **(B)** Spatial distribution of all tumor cells (DAPI), TUNEL-positive cells, bacteria, and collagen in a heterotypic tumor spheroid 36 h after colonization. **(C)** Distribution of normalized fluorescence intensity of collagen, bacteria, and TUNEL as a function of tumor radius. **(D)** Distribution of normalized fluorescence intensity of collagen and bacteria in the zoomed-in region shown in 4A.ii. **(E)** Distribution of the normalized TUNEL signal intensity for the ROI (shown in orange in 4A.i) and the rest of the tumor as a function of tumor radius. Scale bars in A.i and B are 100 μm. The scale bar in A.ii is 10 μm.

To assess the role of bacteria localization on bacteria-induced tumor cell death, we compared the TUNEL signal distribution in the ROI and the rest of the tumor. **Figures 5E** and S6 show a significantly higher signal intensity in ROI compared to the rest of the tumor, indicating a correlation between localized bacterial colonization and regional tumor cell death. These findings underscore the complexity of the interplay between ECM components like collagen, accumulation of bacteria (and other tumor therapeutics), and their antitumor effects.

## Conclusions

Successful bacterial colonization of solid tumors is a requisite for efficacious treatment using BBCT. While recent years have seen much advancement and innovation in engineering new BBCT strains (39), there has been little focus on the important challenge of bacteria interstitial penetration. Using breast cancer tumor models of low and high collagen content, we demonstrate that bacteria penetrate and extensively colonize low collagen-content homotypic tumor spheroids while they remain constrained to the peripheral regions of the physiologically relevant high collagen-content tumor spheroids, even after 48 h. Our mathematical model delineates the role of bacterial growth and diffusivity in the observed colonization patterns and demonstrates that an order of magnitude lower diffusivity in heterotypic tumors dominates the observed outcomes. The reduced bacterial colonization leads to reduced antitumor effects of bacteria. Moreover, quantitative evaluation of the spatial distribution of bacteria, collagen, and apoptotic cells demonstrates a robust correlation between high collagen content, reduced bacterial colonization, and reduced regional tumor cell death. Understanding the effect of collagen on BBCT performance could lead to engineering more efficacious BBCT strains capable of overcoming this barrier to efficacious colonization of primary tumors and metastases.

## Supporting Information

Supporting information is available from

## Acknowledgments

This work was supported by the National Science Foundation (CAREER award, CBET-1454226 to B.B.), Virginia Tech Institute for Critical Technology and Applied Science, and Virginia Tech Center for Engineered Health. Additionally, this work was supported by the National Cancer Institute (R01 CA179556 and CA268069 to S.M.P). The authors express their gratitude to Prof. Aime Franco from Children’s Hospital of Philadelphia for providing the mammary cancer-associated fibroblast cell lines used in this study.

## Author Contributions

BB conceived and supervised the research. YZ, B. Burkel, EJL, SMP, and BB designed the research methodology. YZ conducted the experimental work. EJL performed the mathematical modeling. YZ and B. Burkel carried out the microscopy sample preparation and imaging. All authors contributed to the data analysis. BB led the manuscript preparation. All authors edited the manuscript.

