## Supplementary Information Document for "Tumor Stroma Content Regulates Penetration and Efficacy of Tumor-targeting Bacteria"

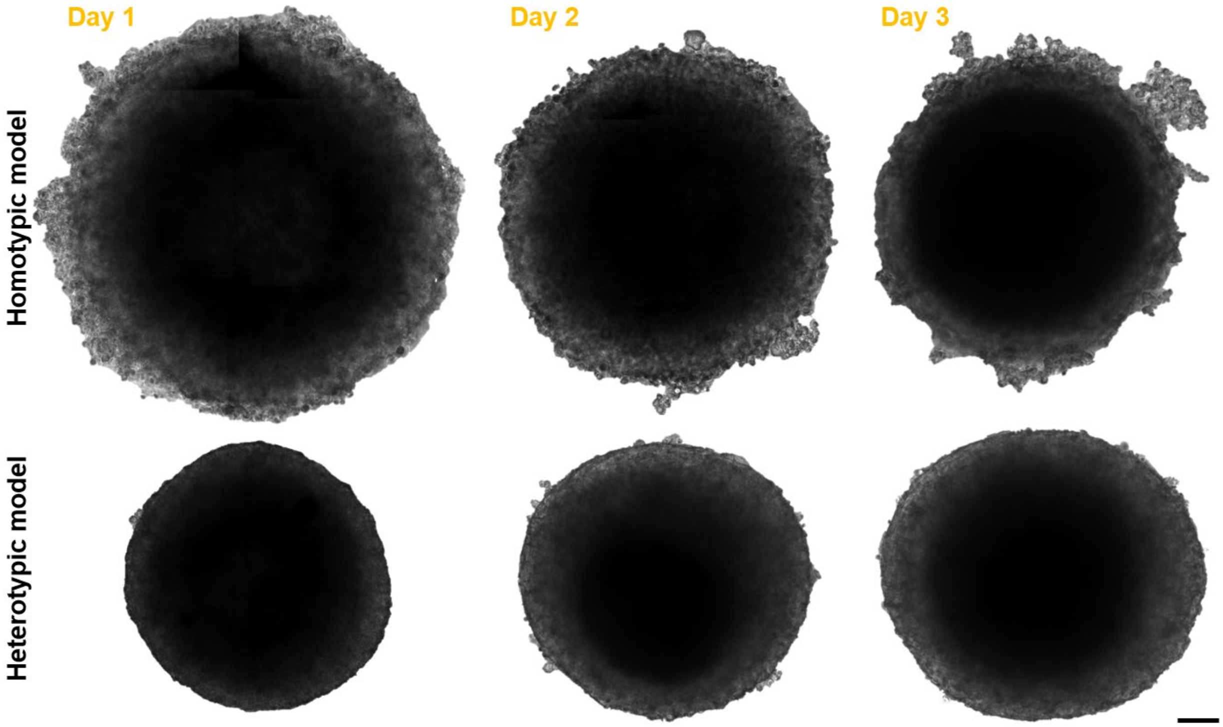


**Figure S1.** Growth of homotypic and heterotypic tumor spheroids. Representative microscopy images of homotypic and heterotypic tumor spheroids in the well plate during the first 3 days. The scale bar is 100 µm.

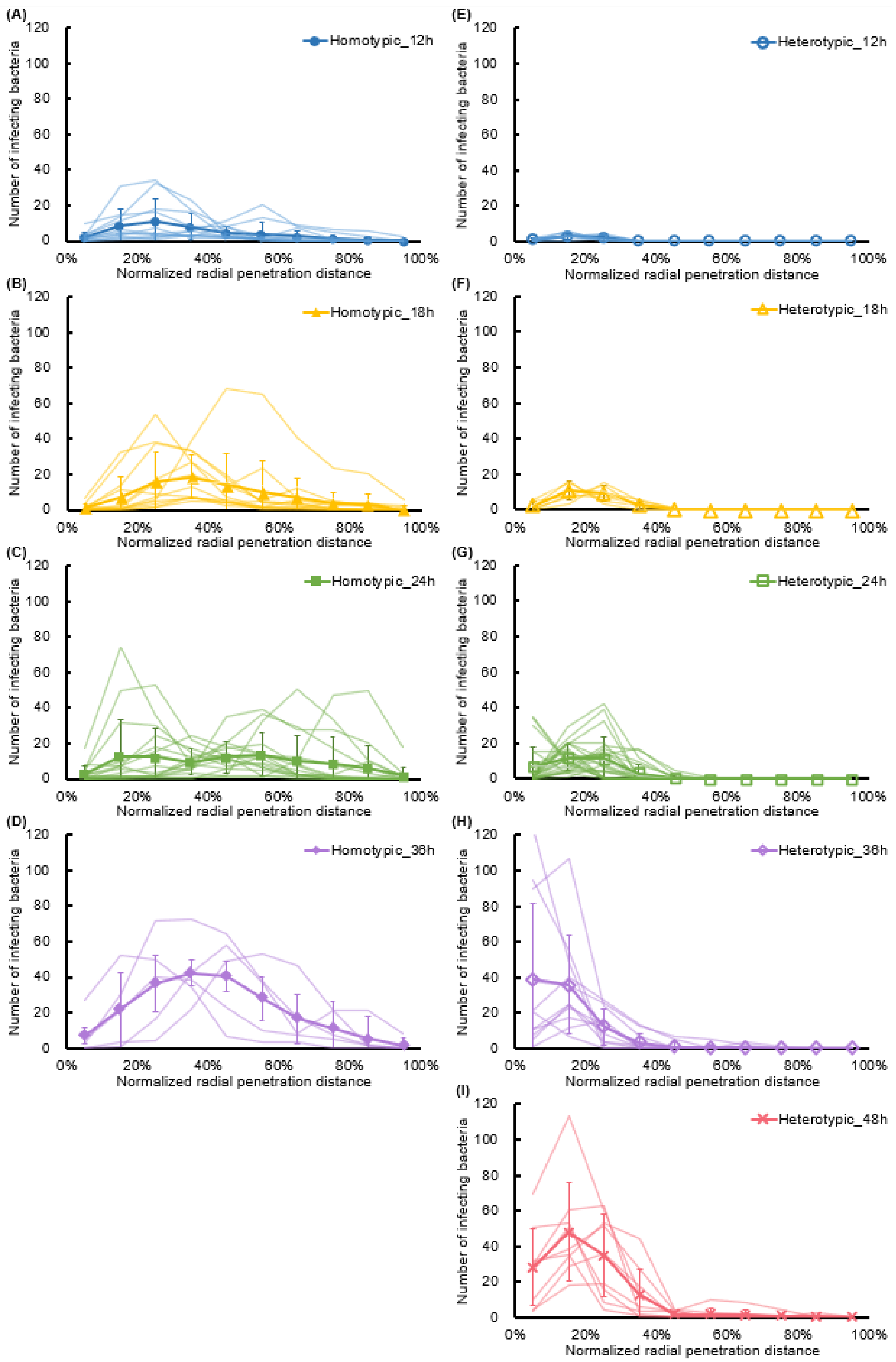


**Figure S2.** Radial distribution of bacteria versus normalized radial penetration distance from the outer boundary in homotypic and heterotypic tumor spheroids. Quantitative analysis of **(A-D)** homotypic and **(E-I)** heterotypic tumor spheroids based on imaged sections at each incubation time. Thin curves show averages across 2-3 sections from each independent tumor spheroid. Thick curves for each case show the averages at each location. Each data point represents the mean ± standard deviation. For homotypic tumor spheroids, n = 11, 13, 16, and 5 independent tumors at 12, 18, 24, and 36 h, and for heterotypic tumor spheroids, n = 8, 7, 20, 11, and 9 independent tumors at 12, 18, 24, 36, and 48 h, respectively.

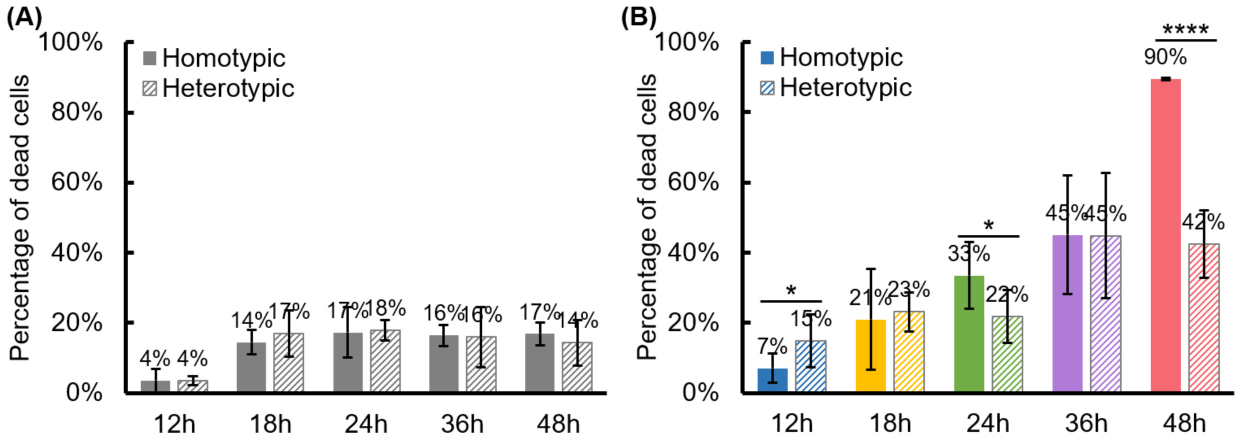


**Figure S3.** Cell viability of homotypic and heterotypic tumor spheroids. Percentage of dead cells of **(A)** control and **(B)** bacteria-treated homotypic and heterotypic tumor spheroids at 12, 18, 24, 36, and 48 h. Each data point represents the mean ± standard deviation. In (A), for control homotypic tumor spheroids, n = 5, 3, 3, 3, and 3 independent tumors, and for control heterotypic tumor spheroids, n = 4, 3, 3, 3, and 3 independent tumors at 12, 18, 24, 36, and 48 h, respectively. In (B), for bacteria-treated homotypic tumor spheroids, n = 10, 3, 8, 5, and 3 independent tumors, and for bacteria-treated heterotypic tumor spheroids, n = 7, 6, 4, 3, and 6 independent tumors at 12, 18, 24, 36, and 48 h, respectively. All results are from the collection of at least 100 total cells from each independent tumor spheroid. Statistical analysis is pairwise comparisons between homotypic and heterotypic tumor spheroids at each time point. * p-value < 0.05, **** p-value < 0.0001.

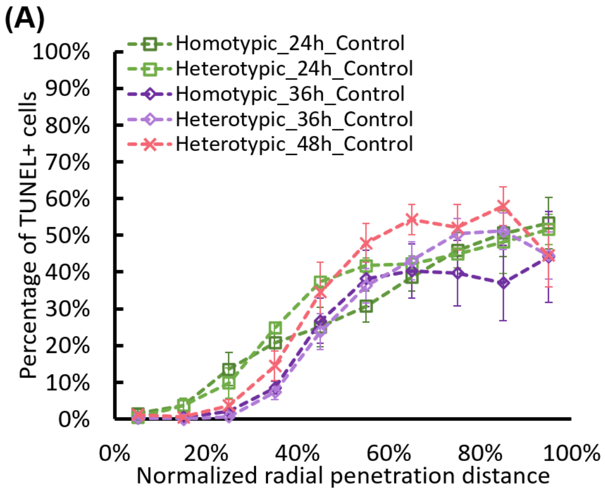


**Figure S4.** Distribution of TUNEL-positive cells in tumor spheroids. **(A)** Distribution of TUNEL-positive cells from the surface to the center of homotypic and heterotypic control tumor spheroids. **(B)** The difference in the average percentage of TUNEL-positive cells between control and bacteria-treated homotypic and heterotypic tumor spheroids. Colored curves for each case show the averages at each timepoint co-plotted. Each data point represents the mean ± standard deviation. For homotypic tumor spheroids, n = 3, 3 (control) and n = 11, 8 (with bacteria), at 24 and 36 h from 3 independent experiments, and for heterotypic tumor spheroids, n = 5, 5, 6 (control) and n = 3, 5, 6 (with bacteria), at 24, 36, and 48 h from 3 independent experiments, respectively.

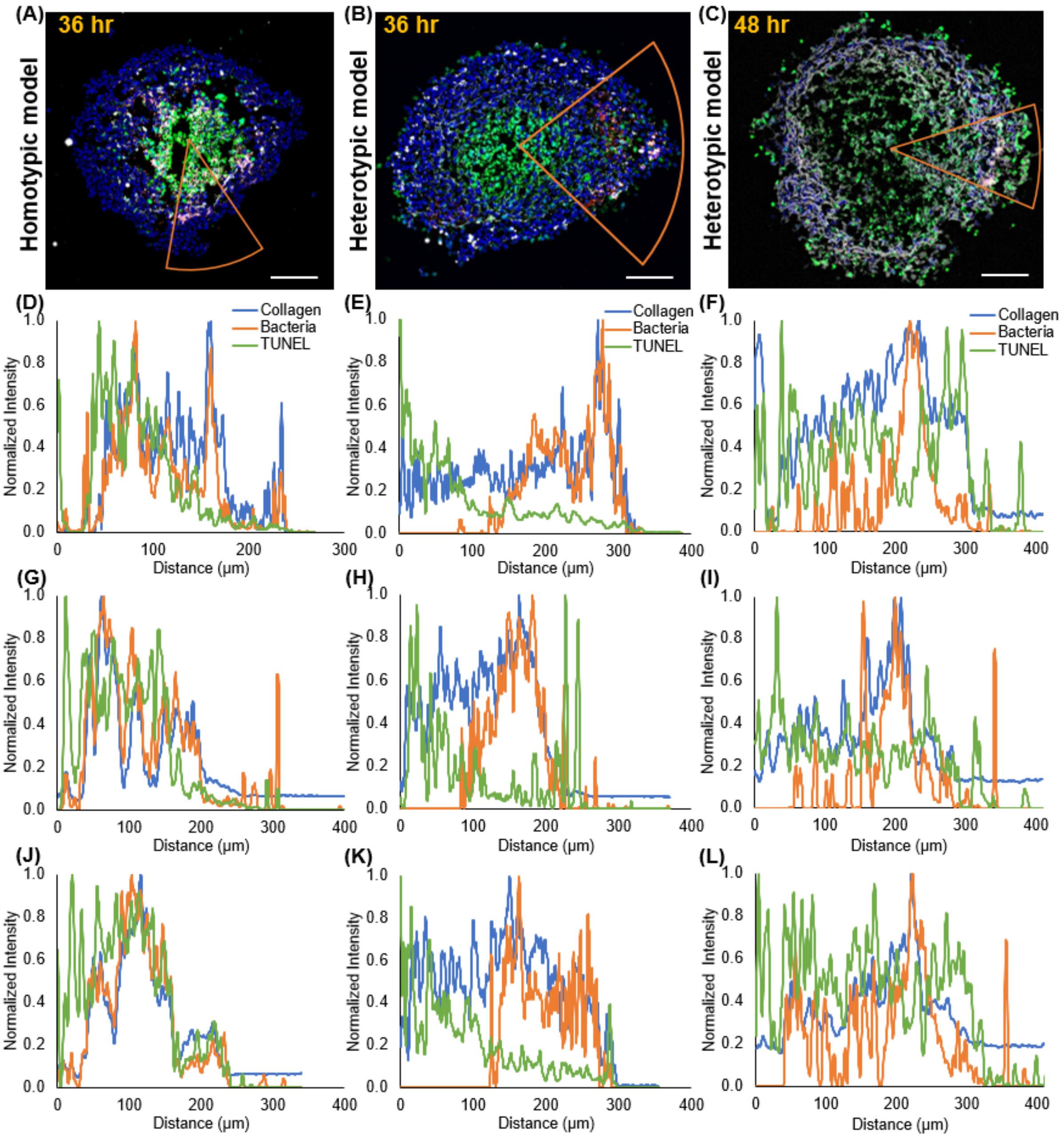


**Figure S5.** Spatial distribution of TUNEL-positive cells, bacteria, and collagen in the homotypic and heterotypic spheroids. Composite fluorescence microscopy image of the spatial distribution of all tumor cells (blue), TUNEL-positive cells (green), bacteria (red), and collagen (white) in **(A)** a homotypic tumor spheroid 36 h after colonization, **(B)** a heterotypic tumor spheroid 36 h after colonization, and **(C)** a heterotypic tumor spheroid 48 h after colonization. **(D-F)** Distribution of normalized fluorescence intensity of collagen, bacteria, and TUNEL-positive cells from the center of the spheroid for the orange slice shown in (A-C). Distribution of normalized fluorescence intensity of collagen, bacteria, and TUNEL-positive cells from the center of the spheroid in high bacterial colonization regions (similar to those shown in (A-C)), for two more homotypic tumor spheroids 36 h after colonization **(G-J)**, two more heterotypic tumor spheroids 36 h after colonization **(H-K)**, and two more heterotypic tumor spheroids 48 h after colonization **(I-L)**. In (A-C), all scale bars are 100 μm.
